# Drug distribution along the cochlea is strongly enhanced by low-frequency round window micro vibrations

**DOI:** 10.1101/2021.05.05.442757

**Authors:** Samuel M. Flaherty, Ian J. Russell, Andrei N. Lukashkin

## Abstract

The cochlea’s inaccessibility and complex nature provide significant challenges to delivering drugs and other agents uniformly, safely and efficiently, along the entire cochlear spiral. Large drug concentration gradients are formed along the cochlea when drugs are administered to the middle ear. This undermines the major goal of attaining therapeutic drug concentration windows along the whole cochlea. Here, utilizing a well-known physiological effect of salicylate, we demonstrate a proof of concept in which drug distribution along the entire cochlea is enhanced applying round window membrane low-frequency micro vibrations with a probe that only partially covers the round window. We provide evidence of enhanced drug influx into the cochlea and cochlear apical drug distribution without breaching cochlear boundaries. It is further suggested that ossicular functionality is not required for the effective drug distribution we report. The novel method of local drug delivery to the cochlea presented here could be implemented when ossicular functionality is absent or impeded and can be incorporated in clinically approved auditory protheses for patients who suffer with conductive, sensorineural or mixed hearing loss.

## Introduction

The relative inaccessibility of the human cochlea and its intricated structure requires new drug delivery technologies to be designed to ensure safe, efficient and uniform drug distribution along the entire cochlear spiral (Salt & Plontke, 2009; Rivera et al., 2012; El Kechai et al., 2015; Hao & Li, 2019). The blood-labyrinth barrier hinders the effectiveness of systemic drug administration to the inner ear (Nyberg et al., 2019) and local drug administration becomes increasingly important. Success of the most frequently used topical, intratympanic drug delivery, when drugs are administrated into the middle ear cavity (Figure 1A), depends on a drugs ability to diffuse into the scala tympani (ST) through the round window membrane (RW) and, to a limited extent, into the scala vestibuli through the oval window occluded by the stapes. If the drug is allowed only to diffuse passively along the narrow, extended ST, its concentration, in theory, should become the same within the entire scala after an arbitrary long time (unrealistic scenario, Figure 1B) (Sadreev et al., 2019). However, for a drug to be effective, it has to be cleared from the ST into other cochlear compartments (more realistic scenario, Figure 1B). Dynamic equilibrium between diffusion and clearing leads to the formation of substantial steady-state, base-to-apex drug concentration gradients along the cochlea (Salt & Ma, 2001; Sadreev et a., 2019), which have been confirmed experimentally for marker ions and contrasting agents (Salt & Ma, 2001; Haghpanahi et al., 2013), corticosteroids (Plontke et al., 2008; Creber et al., 2018) and antibiotics (Mynatt et al., 2006; Plontke et al., 2007). Thus, intratympanic drug administration faces the fundamental problem of limited passive diffusion within the cochlea, which undermines drug efficiency due to the inability of drugs to reach their targets within the therapeutic concentration window.

**Figure 1.**
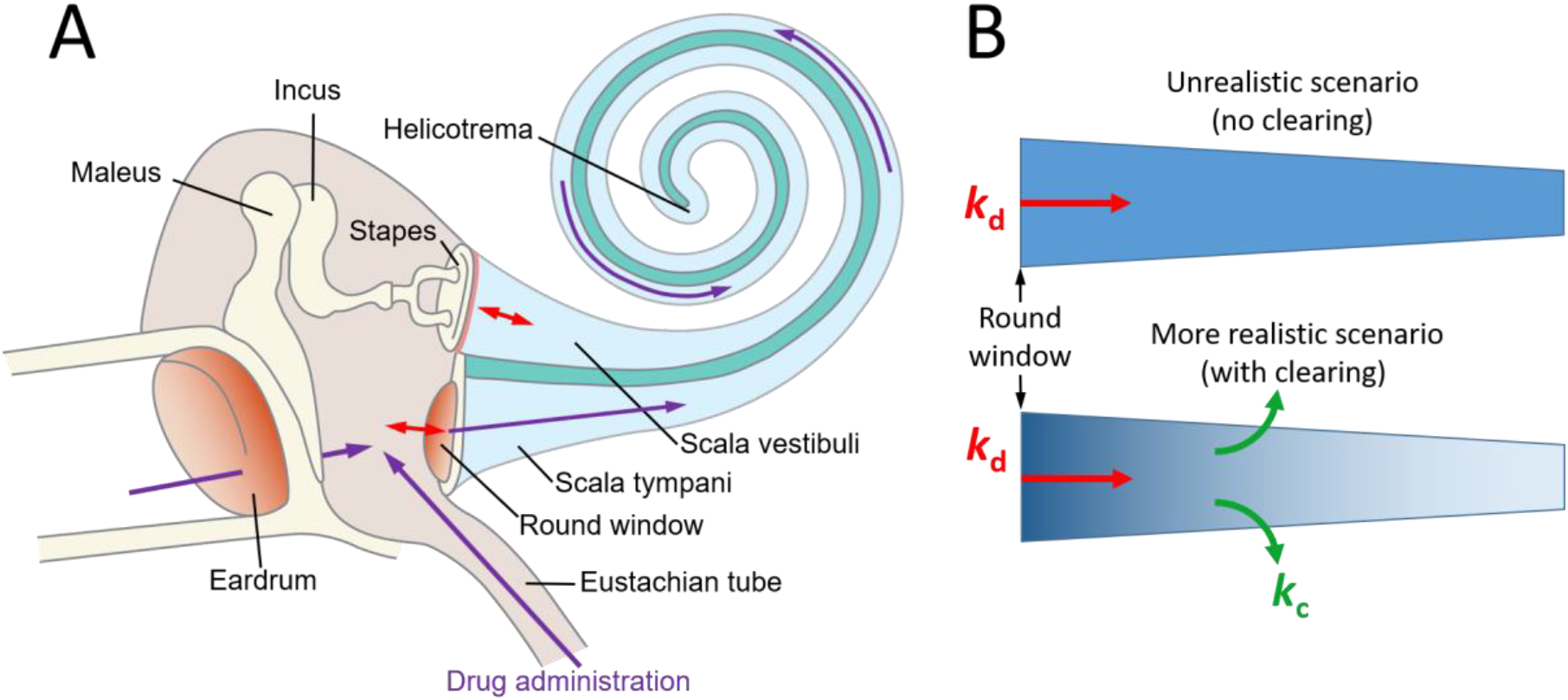
Schematic of the mammalian hearing organ (**A**) and two scenarios of molecular drug diffusion along the scala tympani (**B**). (**B**) Passive molecular diffusion of a drug along the scala tympani is described by a diffusion (*k*_d_) and clearing (*k*_c_) coefficients. For a given geometry of the scala tympani, the steady-state drug concentration gradient (denoted by the blue colour intensity) along it depends only on the ratio *k*_d_/*k*_c_ (Sadreev et al., 2019). (**A**) is modified from (Lukashkin et al., 2020).

A few relatively non-invasive techniques for assisting substance mixing along the cochlea have been suggested recently that utilise low-frequency pressure stimulation (Lukashkin et al., 2020), stimulation at acoustic frequencies (Park & Moon, 2014; Shokrian et al., 2020) and ultrasound (Liao et al., 2020) which cause reciprocated movement of the stapes and RW. While the later method relies on the formation of ultrasound-induced microbubbles which can act directly on the RW (Liao et al., 2020), the other techniques require the mobility of the ossicular chain. If the ossicular chain is immobile or malformed then these techniques become non-applicable. In this study we demonstrate that, in this case, micro vibrations of the RW alone can facilitate drug distribution along the cochlear spiral.

## Materials and methods

### Animals and surgery

Animal preparation and signal generation and recording have been described elsewhere (Burwood et al., 2017). Briefly, pigmented guinea pigs of similar weight (350-360 g) and both sexes were anaesthetised with the neurolept anaesthetic technique (0.06 mg/kg body weight atropine sulphate s.c., 30 mg/kg pentobarbitone i.p., 500 μl/kg Hypnorm i.m.). Additional injections of Hypnorm were given every 40 minutes. Additional doses of pentobarbitone were administered as needed to maintain a non-reflexive state. The heart rate was monitored with a pair of skin electrodes placed on both sides of the thorax. The animals were tracheotomized and artificially respired with a mixture of O_2_/CO_2_, and their core temperature was maintained at 38°C with a heating blanket and a heated head holder.

All procedures involving animals were performed in accordance with UK Home Office regulations with approval from the University of Brighton Animal Welfare and Ethical Review Body.

### Signal generation and recording

The middle ear cavity of the ear used for the measurements and salicylate application was opened to reveal the RW. Compound action potentials (CAPs) of the auditory nerve in response to pure tone stimulation were measured from the cochlear bony ridge in the proximity of the RW membrane using Teflon-coated silver wire coupled to laboratory designed and built extracellular amplifier (James Hartley). Thresholds of the N1 peak of the CAP at different frequencies, which corresponds to different distances from the cochlear base (Greenwood, 1990), were estimated visually using 10 ms pure tone stimuli at a repetition rate of 10 Hz.

For acoustic stimulation sound was delivered to the tympanic membrane by a closed acoustic system comprising two Bruel and Kjaer 4134 ½” microphones for delivering tones and a single Bruel and Kjaer 4133 ½” microphone for monitoring sound pressure at the tympanum. The microphones were coupled to the ear canal via 1 cm long, 4 mm diameter tubes to a conical speculum, the 1 mm diameter opening of which was placed about 1 mm from the tympanum. The speculum was sealed in the ear canal. The closed sound system was calibrated in situ for frequencies between 1 and 50 kHz. Known sound pressure levels were expressed in dB SPL re 2×10^−5^ Pa.

All acoustic stimuli in this work were shaped with raised cosines of 0.5 ms duration at the beginning and at the end of stimulation. White noise for acoustical calibration and tone sequences for auditory stimulation were synthesised by a Data Translation 3010 board (Measurement Computing Corporation, MA) at 250 kHz and delivered to the microphones through low-pass filters (100 kHz cut-off frequency). Signals from the acoustic measuring amplifier (James Hartley) were digitised at 250 kHz using the same board and averaged in the time domain. Experimental control, data acquisition and data analysis were performed using a PC with custom programmes written in MATLAB (MathWorks, MA).

### Salicylate application

5 μl of 100 mM sodium salicylate solution in Hanks’ Balanced Salt Solution were placed on the RW using pipettes. The solution was removed from the RW using paper wicks to observe the wash out effect.

### Round window stimulation

A miniature loudspeaker K16-50 Ohm (Visaton GmbH, Haan, Germany) was used to vibrate the RW. Loudspeaker dust cover was removed and a carbon rod (20 mm in length and 0.5 mm in diameter) was glued centrally on the loudspeaker membrane. The probe was perpendicular to the loudspeaker face and remained in this position during experiments to ensure no sideward movements. The probe tip was rounded using a thin layer of superglue preventing RW damage and carbon rod fragmentation. The loudspeaker was fixed to a steel rod, using araldite, and the rod was held in a micromanipulator for a precise probe placement. A programmable synthesiser/signal generator (Philips PM5193) was used to drive the loudspeaker in the experiments. The probe movements versus voltage applied to the loudspeaker were calibrated prior to experiment by focusing a laser vibrometer (CLV-2534, Polytec GmbH, Waldbronn, Germany) at the probe tip along the probe axis and measuring dependence of the probe vibration velocity on the voltage applied to the loudspeaker at the RW stimulation frequencies. The probe vibration amplitude was calculated by integrating its velocity.

During experiments, the carbon probe was placed at about 45-degrees to the RW because of a limited access to the RW. Probe vibrations started immediately after placing salicylate solution on the RW. In the first 20 minutes period, acoustic CAP threshold recordings were taken every 3-5 minutes to record the fast action of salicylate at the basal region of the cochlea. Due to the very low frequencies used to vibrate the RW, there was no CAP generated in response to the probe vibrations, allowing recordings of the CAP due to acoustic stimulation to be taken during the RW micro vibrations. After 20 minutes, the CAP threshold recordings were made every 10 minutes until a total of 60 minutes of RW micro vibrations. To washout, the carbon probe was removed, the salicylate was removed from the RW using fine paper wicks and the recovery of CAP threshold to acoustic stimulation was recorded.

### Recording of stapes vibrations

Stapes vibrations were recorded using a laser vibrometer (CLV-2534, Polytec GmbH, Waldbronn, Germany). The laser beam was focussed on the stapes head. The output voltage from the vibrometer was low-pass filtered at 100 kHz, with a sensitivity of 2 mm/s/V.

### Fluorescent dye experiments

Lucifer yellow CH, lithium salt (Thermo Fisher Scientific) was used to visualize diffusion in straight water filled pipes. The pipes with an approximate length of 40 mm were constructed using Tygon™ LMT-55 tubing (1.14 mm ID, 0.80 mm wall, Fisher Scientific). An outlet was made with a 25G needle and inserted through the pipe’s wall close to one end and fixed in place with superglue. A membrane, cut from a laboratory latex glove (typical thickness of 0.1 mm), was glued with superglue at the same pipe end making sure that the glue does not cover the open surface of the membrane. The other pipe end was closed with a Blu Tack (Blue-tack.co.uk) plug to prevent water evaporation. A 25G needle was inserted through the plug into the pipe to provide pressure relief. The outlet was used to fill the pipe with deionized water to a distance of about 30 mm from the latex membrane and to inject 0.2 μl of 5% Lucifer yellow water solution into the pipe using a pipette. Lucifer yellow fluorescence was excited using a 470 nm laser source (Dragon Lasers, Changchun Jilin, China) and still images were taken (Sony α6100 camera, Sony Macro E 30mm F/3.5 lens) through an optical band pass filter (FB540-10, Thorlabs Inc.) to assess dye diffusion over time. The same miniature loudspeaker K16-50 Ohm (Visaton GmbH, Haan, Germany) with the carbon probe attached as used for the RW stimulation was employed to vibrate the latex membrane in assisted diffusion experiments. The carbon probe touching the membrane was pushed slightly toward inside of the pipes at rest to ensure membrane tension and its relaxation during backward phase of probe strokes. Fluorescence intensity profiles were measured along the pipe axis using Fiji open source image processing package.

## Results

### Low-frequency round window membrane micro vibrations do not elevate hearing thresholds in guinea pigs

RW stimulation with the carbon probe did not evoke any electrical responses that could be detected by the RW electrode, which made it possible to make continuous CAP threshold measurements to acoustic stimulation throughout the probe vibrations. Hearing sensitivity, assessed by measured CAP thresholds, did not change when 10 μm peak-to-peak continuous probe vibrations were applied to the RWM at 2 or 4 Hz for up to 60 minutes (Figure 2). In our experiments, the probe covered only a small part of the RW. Under these conditions, most of the pressure relief during the probe movement is through the RW area not occluded by the probe. The generated far-field pressure component is small and cochlear excitation is due mainly due to RW near-field pressure, which excites a conventional travelling wave at acoustic frequencies (Weddell et al., 2014). Had a significant far-field pressure been generated, it would cause a stapes movement. However, we were not able to detect any stapes responses above the measurement noise floor of ~ 0.1 nm during the RW probe vibrations either at 2 or 4 Hz. As indicated by measurements from the RW electrode, the near-field pressure did not excite the cochlear sensory apparatus at the frequencies of 2 and 4 Hz used in our experiments, even for relatively large 10 μm RW probe displacement. A consequence of this finding is that even large probe induced vibrations of the RW membrane at these frequencies should be safe and unlikely to produce hearing loss (see Discussion for detailed analysis).

**Figure 2.**
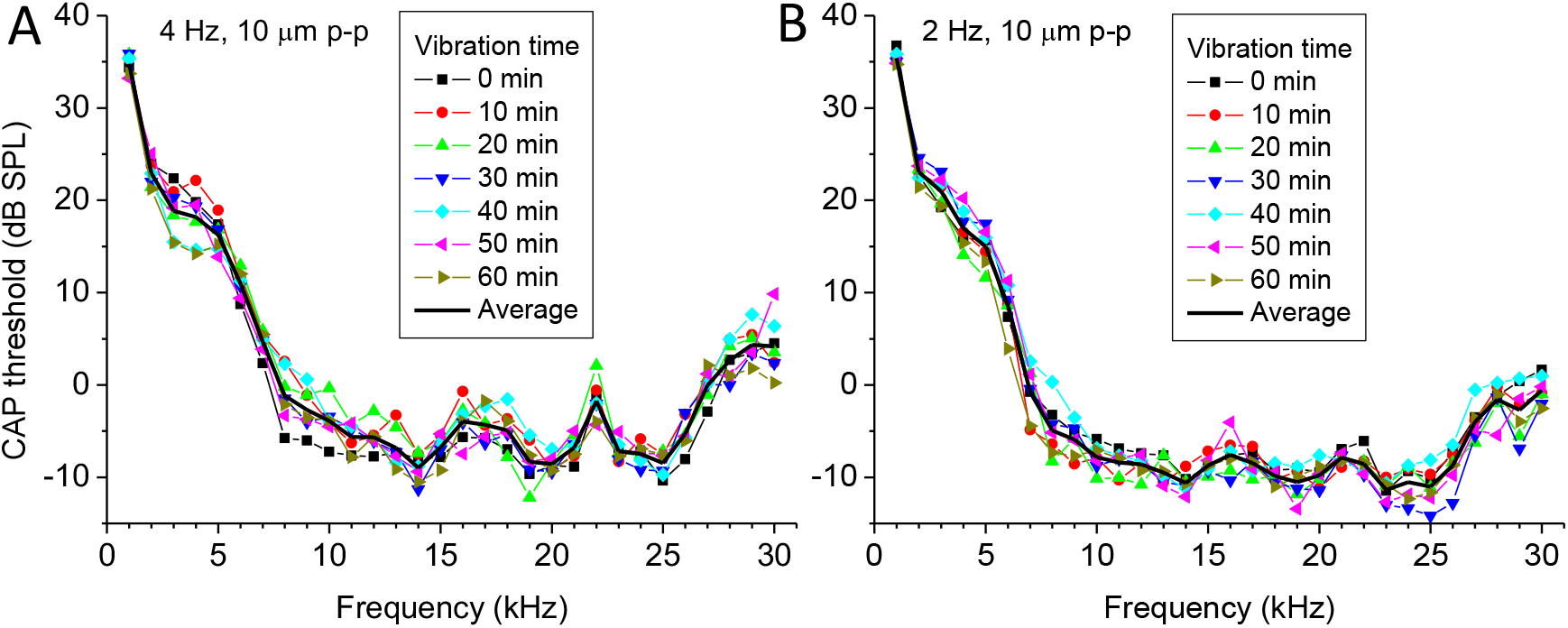
The effect of continuous RW probe vibrations at frequencies of 4 (**A**) and 2 (**B**) Hz on acoustic CAP thresholds without application of salicylate solution as a function of acoustic stimulus frequency. Frequency and amplitude of RW probe vibrations is indicated for each panel. Corresponding duration of vibrations is indicated by curves with different symbols. Each curve represents averaged data for 4 preparations (mean value, SD is not indicated for clarity). Solid black curves indicate averaged data (mean value) for all times presented at each panel. Vibration time indicated corresponds to the beginning of each individual CAP threshold curve measurements. It took less than a minute to record the CAP threshold curve for the entire frequency range 1-30 kHz.

### Round window membrane micro vibrations promote drug distribution along the cochlear spiral

The ability of micro vibrations of the RW to improve drug distribution along the cochlear spiral was demonstrated in our experiments with the application of salicylate to the RW. Salicylate readily diffuses through the RW (Borkholder et al., 2014; Sadreev et al., 2019). To monitor salicylate diffusion along an intact guinea pig cochlea *in vivo*, we utilized the suppressive effect of salicylate on cochlear amplification by blocking the outer hair cell (OHC) somatic motility (Russell and Schauz, 1995; Hallworth, 1997). Salicylate competitively binds the motor protein prestin, essential for OHC motility, by repelling Cl^−^ -ions and preventing interaction with the anion-binding site (Oliver et al., 2001). We measured the elevation of CAP thresholds caused by salicylate at different frequencies of acoustic stimulation, which, due to cochlear tonotopicity, corresponds to different distances from the RW (Greenwood, 1990). Thus, through measuring the CAP threshold elevations we could assess the spread of salicylate along the cochlea when it was applied to the RW.

When 5 μl of 100 mM/l salicylate solution was applied to the RW (Figure 3), it caused a rapid increase followed by saturation of CAP thresholds for high frequency tones with the characteristic frequency place situated below or close to the RW (e.g. 25 kHz, Figure 3B, D). Over time, CAP threshold elevation gradually spreads to lower frequencies (Figure 3A, C) indicating salicylate diffusion into the cochlear apex. Salicylate did not cause elevation of the CAP threshold responses for frequencies below 5 kHz, which corresponds to about 45% of the total cochlear length from the base, when it diffused through the cochlea passively (Sadreev et al., 2019). The calculated gradient of base-to-apex salicylate concentration was about 13 orders of magnitude. When, however, placement of salicylate solution on the RW was followed by RW probe vibrations at frequencies of 2 and 4 Hz, the CAP threshold was elevated throughout the entire 1–30 kHz frequency range tested (Figure 3). The CAP threshold elevation did not saturate and was still rising for frequencies below 5 kHz (Figure 3B, D) indicating continuous increase in salicylate concentration in this cochlear region even after 60 minutes of the probe vibration. This corresponds to about 25% of the total cochlear length from the apex (Greenwood, 1990). Partial recovery of the CAP thresholds during washing out salicylate from the RW after 60 minutes of its application (Figure 3) provided confirmation that the integrity of the sensory cells was preserved and the CAP threshold elevation after joint salicylate application and RW probe vibrations was not caused by the later.

**Figure 3.**
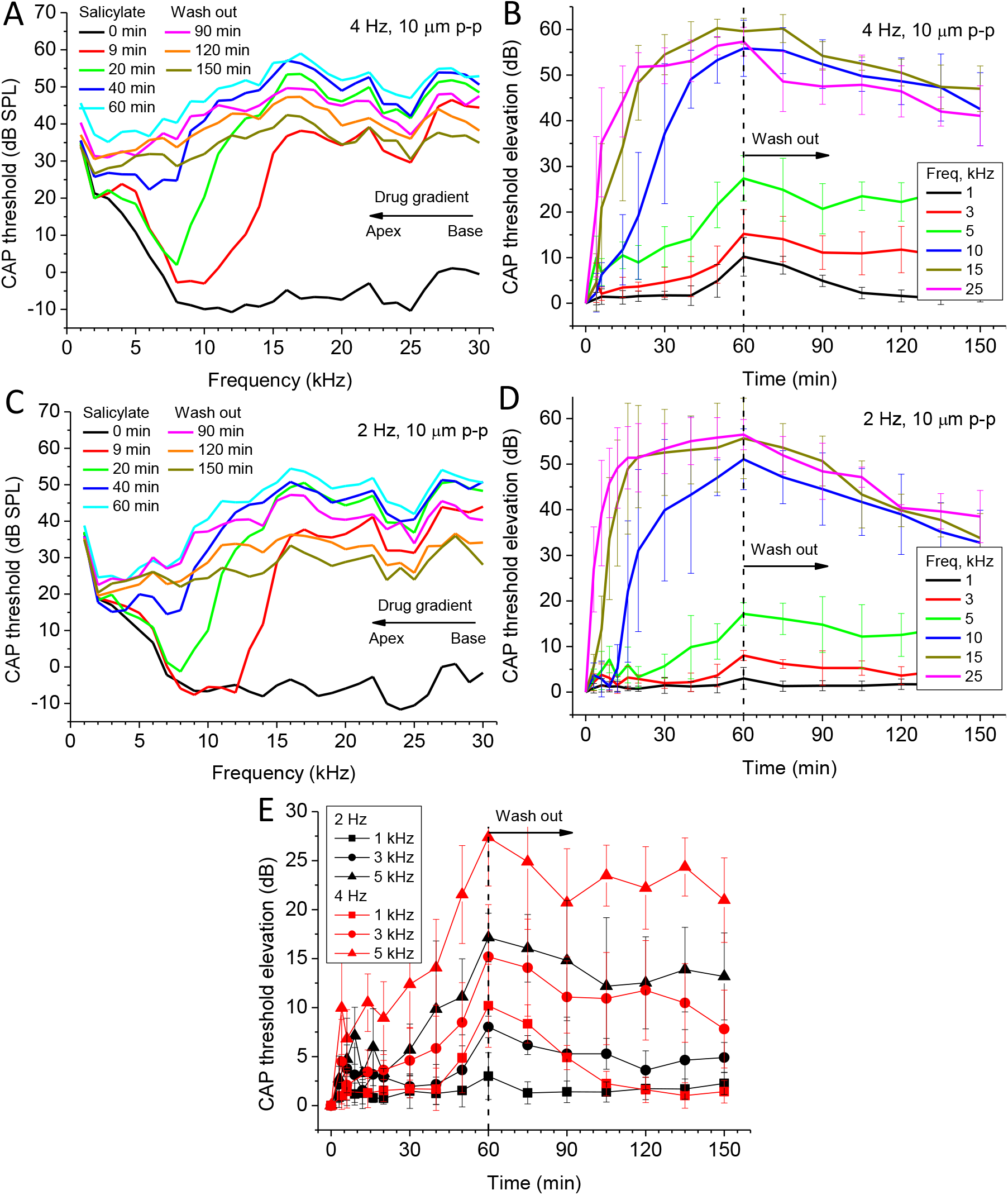
Effect of combined application of 5 μl 100 mM/l salicylate solution and continuous RW probe vibrations at frequencies of 4 Hz (**A, B, E**) and 2 Hz (**C, D, E**) as a function of acoustic stimulus frequency. Salicylate was applied at time zero and RWM probe vibrations started at the same time. Salicylate was washed out after 60 minutes. (**A, C**). CAP thresholds for different times of salicylate application/RW vibrations (colour coded curves) as a function of acoustic stimulus frequency (mean values, SDs are not shown for clarity, N = 4). (**B, D**). CAP threshold elevations relative to the thresholds before salicylate application (time zero) for a few acoustic stimulus frequencies (colour coded curves) which correspond to different locations along the cochlea (mean ± SD, N = 6 and 4 for (**B**) and (**D**) respectively). (**E**). CAP threshold elevations relative to the thresholds before salicylate application (time zero) for acoustic stimulus frequencies (different symbols) corresponding to apical half of the cochlea (mean ± SD, N = 6 and 4 for 4 and 2 Hz of probe vibrations respectively). Statistically significant (p < 0.05, unpaired *t*-test) differences between the threshold elevations for probe vibrations at 4 and 2 Hz are observed after 60 minutes of salicylate application/RW vibrations.

### Drug distribution along the cochlea length depends on the frequency of round window micro vibration

During combined application of salicylate solution to the RW and RW probe vibrations, the CAP threshold elevations increase when the frequency of the RW probe vibrations is increased (Figure 3). This is particularly evident for the lowest frequencies of acoustic stimulation (Figure 3E). For the same acoustic frequencies (i.e. cochlear locations), the total CAP threshold elevations after 60 minutes of combined salicylate application and RW probe vibrations at 4 Hz were significantly higher (p < 0.05, unpaired *t*-test) than the threshold elevations observed during probe vibrations at 2 Hz. This frequency dependence confirms that increase in the CAP thresholds at frequencies which correspond to more apical cochlear locations and, hence, enhanced diffusion of salicylate to the cochlear apex, was not due to placement of the probe alone and probe vibrations were required to observe the effect.

### Comparison between different techniques of drug delivery through the round window membrane

When 5 μl of 100 mM/l salicylate solution was applied to the RW and salicylate is allowed to diffuse passively along the cochlear, it does not cause the CAP threshold elevations for frequencies of acoustic stimulation below 5 kHz (Figure 4, black line; Sadreev et al., 2019). This effectively means that salicylate is not able to reach the apical 50% of the cochlea at any effective concentrations. Combined application of salicylate and low-frequency (4 Hz) pressure oscillations to the ear canal (cochlear pumping), which causes large amplitude (80 μm peak-to-peak), movement of the stapes and reciprocal movement of the RW, causes elevation of CAP thresholds within the entire frequency range tested (Figure 4, blue line; Lukashkin et al., 2020). This indicates the ability of the cochlear pumping to distribute salicylate evenly along the entire cochlea. Joint application of salicylate and RW probe vibrations (4 Hz, 10 μm peak-to-peak amplitude) reported in this study causes CAP threshold elevations for the frequencies corresponding to the cochlear apex (Figure 4, red line) indicating enhanced drug diffusion during the RW vibration. These threshold elevations at the cochlear apex (below 5 kHz) were smaller than those observed during cochlear pumping, probably because of the smaller RW probe vibration amplitude (10 μm peak-to-peak) compared to the stapes vibration amplitude (80 μm peak-to-peak). However, the CAP threshold elevations for the basal half of cochlea (frequency of acoustic stimulation above 5 kHz) observed during the RW probe vibrations exceed those recorded during both passive salicylate diffusion and cochlear pumping. These high-frequency thresholds elevations during the RW vibrations are, in fact, close to the maximum threshold elevations after complete block of the cochlear amplifier by application of 1 M/l salicylate solution (Figure 4, circles; Sadreev et al., 2019) and indicate higher basal concentrations, i.e. influx of salicylate into the ST. Therefore, the RW probe vibrations not only promote drug diffusion into the cochlea apex but also enhance salicylate passage through the RW.

**Figure 4.**
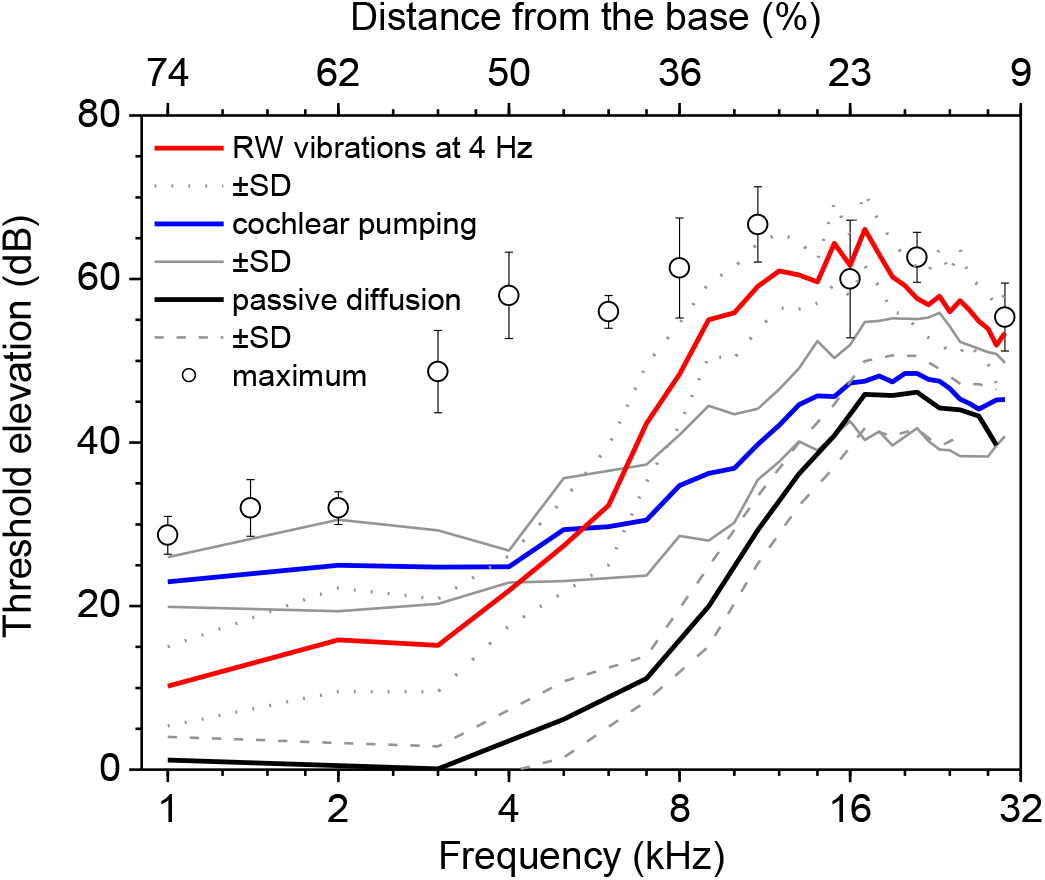
Comparison between different techniques of drug delivery through the RW. Frequency dependence of the CAP threshold elevation after 60 min of 5μl 100 mM/l salicylate solution application for passive diffusion (black line, mean ± SD, N = 5, Lukashkin et al., 2020), during cochlear pumping (35 min of the total pumping time) (blue line, mean ± SD, N = 5, Lukashkin et al., 2020) and continuous RW probe vibrations at 4 Hz (red line, mean ± SD, N = 6) are shown. Open circles show maximal increase of the CAP thresholds after complete block of the cochlear amplifier by application of 5 μl of 1 M/l salicylate solution to the RW (mean ± SD, N = 3) (Sadreev et al., 2019).

### Passive and assisted diffusion of fluorescent dye

To gain insight into the mechanism of enhanced drug diffusion in our experiments with the RW vibrations, we compare the speed of passive, molecular diffusion of Lucifer yellow along water filled straight pipes and its diffusion assisted by micro vibrations of a membrane covering one end of the pipes (Figure 5A, B). The pipes were water filled to ~ 30 mm from the membrane and their internal cross-sectional area was ~ 1.02 mm^2^ which correspond to the length and average cross-sectional area of the human ST, respectively (Thorne et al., 1999). Fluorescence intensity profiles, obtained immediately after injection of 0.2 μl of the dye, were closely similar in all experiments (Figure 5C), indicating the initial conditions remained constant for all sets of measurements. Small, 40 μm peak-to-peak vibration of the membrane at 10 Hz over 60 minutes enhanced dye distribution compared to passive diffusion (Figure 5D). The additional spread of the diffusion front was small (about 1.7 mm for fluorescence intensity of 150 AU, green arrow in Figure 5D). However, due to nonlinear dispersion of the diffusion front, this small additional spread led to a statistically significant increase (unpaired *t*-test for 0.5 mm bins, p<0.05) in the fluorescence intensity, i.e. dye concentration, over a much wider range of 9 mm (blue horizontal bar, Figure 5D).

**Figure 5.**
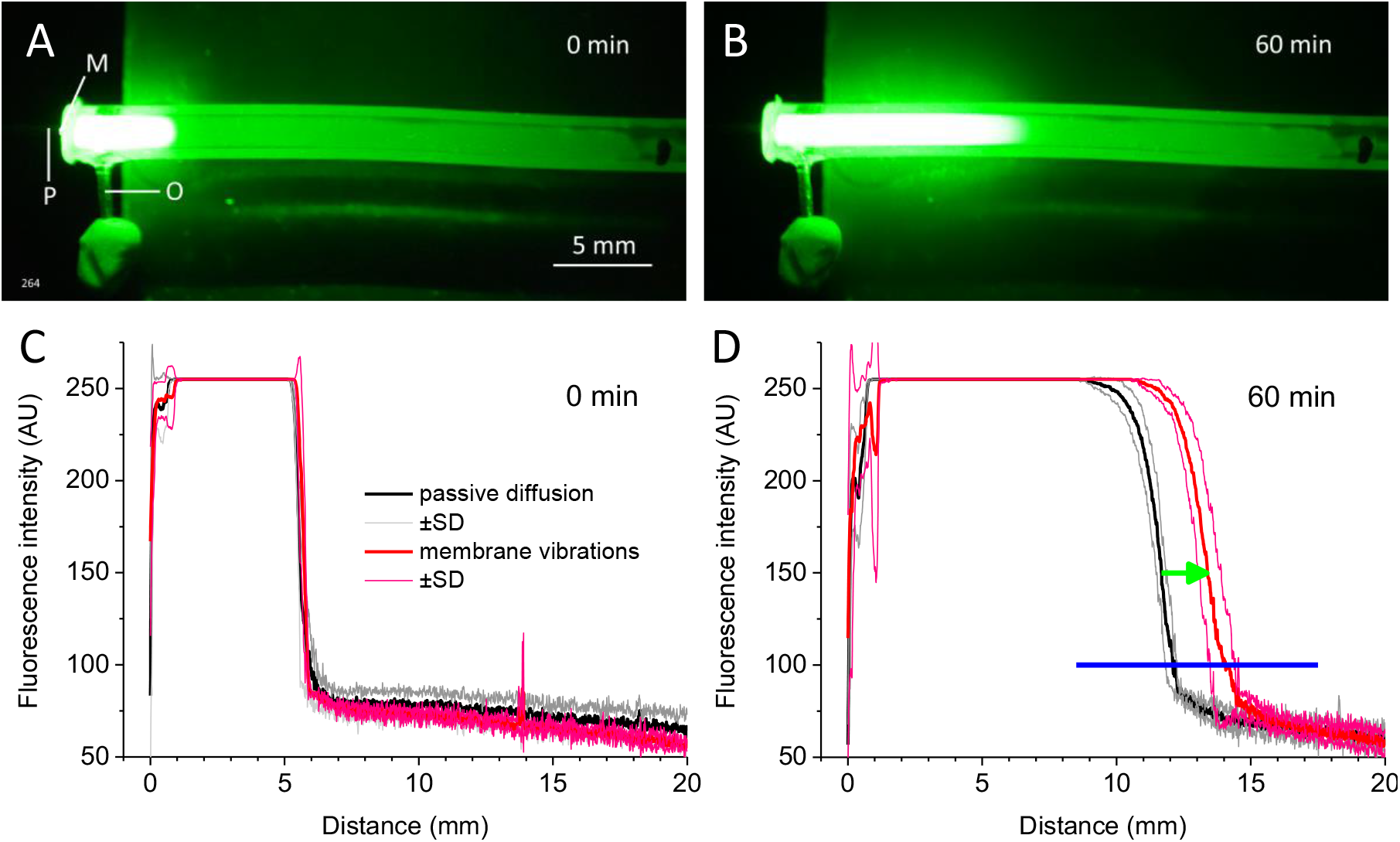
Distribution of Lucifer yellow in a straight pipe during passive diffusion and during vibrations of a membrane at a pipe end. (**A**). Fluorescence immediately after injections of 0.2 μl of 5% Lucifer yellow water solution into the pipe through outlet O. M – latex membrane; P – carbon probe. (**B**). Fluorescence after 60 minutes of membrane vibrations at 10 Hz with 40 μm peak-to-peak carbon probe movements. (**C**). Overlapping fluorescence intensity profiles measured along the pipe axis for all passive diffusion (black line, mean ± SD, N = 3) and membrane vibration (red line, mean ± SD, N = 3) experiments indicating the same initial conditions immediately after the dye injection. (**D**). Fluorescence intensity profiles after 60 minutes of passive dye diffusion (black line, mean ± SD, N = 3) and membrane vibrations (red line, mean ± SD, N = 3). The same experiments as in C. Green arrow indicates additional spread of the diffusion front during membrane vibrations. Blue horizontal bar indicates the spread of statistically significant increase (unpaired *t*-test for 0.5 mm bins, p<0.05) in the fluorescence intensity/dye concentration which is observed during membrane vibrations.

## Discussion

This is a proof of concept report which demonstrated that vibrating the partially occluded RW at low frequencies of 2 and 4 Hz and with an amplitude of 5 μm facilitates drug distribution along the cochlear spiral. Finding optimal and safe parameters of the RW vibrations was outside the study’s scope. However, we can conclude that for the range of stimulation parameters used within the timeframe of experiments, drug diffusion enhances with increasing the RW stimulation frequency without affecting neural thresholds. This frequency dependence of the drug distribution also indicates that placement of the RW probe did not affect the RW integrity. At the same time, enhanced effect of the RW vibrations on the CAP threshold elevation in the basal half of cochlea (Figure 3) compared to passive diffusion (Sadreev et al., 2019) and cochlear pumping (Lukashkin et al., 2020) suggests that the RW drug permeability for salicylate was increased during direct mechanical stimulation of the RW.

RW vibration stimulation alone did not elicit RW electrical responses, including CAPs associated with afferent fibre/inner hair cell excitation or cochlear microphonic potentials dominated by basal turn OHC mechanoelectrical transducer currents (Patuzzi et al., 1989; Cheatham et al., 2011). Previously, very large cochlear microphonic potentials in response to 5 Hz *acoustic* stimulation were recorded from the cochlear apex but not the cochlear base in guinea pigs suggesting excitation of the OHCs in this tonotopic frequency place (Salt et al., 2013). We argue, however, that excitation of sensory hair cells due to the low-frequency RW vibrations was minimal in our experimental configuration. The RW probe diameter (0.5 mm) and its cross-sectional area (0.2 mm^2^) were much smaller than the dimensions and area of the RW in guinea pigs (Ghiz et al., 2001; Wysocki et al., 2005), and the probe covered only a small part of the RW. Under these conditions, most of the pressure relief during the probe movement was through the RW area not occluded by the probe (Weddell et al., 2014) and the average alternating far-field pressure, *P*_M_, generated within the cochlea was small. This was confirmed by the absence of stapes responses, which were below the measurement noise floor (~ 0.1 nm) during the probe vibrations at RW either at 2 or 4 Hz. The magnitude of *P*_M_ will depend on the stiffness, *S*, of the freely moving area, *A*, of the RW and its volume velocity, *q*, as (Weddell et al., 2014)

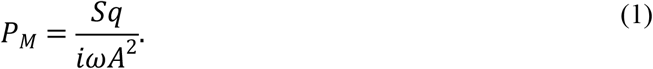

Even if a small far-field pressure was generated in our experiments due to finite stiffness, *S*, of the RW, which did not generate measurable stapes vibrations, then it still would not lead to a significant excitation of the BM. Frequencies of 2 and 4 Hz used in our experiments are notably below the helicotrema cut-off frequency in guinea pigs (Marquardt et al., 2007) and will be filtered out by the helicotrema, preventing the BM excitation at these frequencies and damage to the cochlear sensory cells during the RW probe stimulation.

Vibrations of the partially occluded RW at acoustic frequencies excite the basilar membrane with conventional travelling waves. A jet-like, near-field component, *P*_N_, of a complex pressure field near the RW is the proposed mechanism of stimulation (Weddell et al., 2014). *P*_N_ is proportional to the fluid density, *ρ*, and to the acceleration of the probe, *iωq*, and an indicative overall magnitude of *P*_N_ can then be defined as

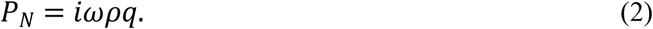

Because RW stimulation with the carbon probe did not evoke any cochlear microphonic potentials from the basal OHCs that could be detected by the RW electrode (Patuzzi et al., 1989; Cheatham et al., 2011), we conclude that this near-field component did not excite the basilar membrane at frequencies of 2 and 4 Hz used in our experiments which also resulted in lack of excitation of the cochlear sensory apparatus and absence of any probe induced hearing loss even for relatively large 10 μm peak-to-peak RW probe displacements. It is worth noting, that the near-field pressure, *P*_N_, increases with increasing frequency (Equation 2). This can explain the higher efficiency of 4 Hz RW stimulation compared to 2 Hz (Figure 3E) if the near-field pressure component is the main factor facilitating enhanced drug diffusion during vibration of a partially occluded RW.

The question is how this short acting jet-like, near-field pressure component can facilitate drug distribution along the entire cochlea which is an order of magnitude longer than the near-field pressure spread (Weddell et al., 2014). The fluorescent dye experiments (Figure 5), while being different from the RW stimulation experiments in two important aspects, provide an insight into the underlying physical mechanisms. Firstly, the latex membrane stiffness was much larger than the RW stiffness. Pressure relief in this case was through the open pipe end and a large far-field pressure component was generated within the fluid-filled pipe leading to movement of the entire fluid column. Taylor dispersion (Taylor, 1953) of solvents is observed during oscillatory pipe flows which lead to additional spread of solvents compared to molecular diffusion alone (Aris, 1960; Watson, 1983). It has been demonstrated experimentally that for small-stroke fluid oscillatory movements and dimensions of the human cochlea this effect is small (Dasgupta, 2015), which is confirmed by lack of changes in the diffusion front in our experiments (Figure 5D). However, when a physical body vibrates in confined spaces, which resembles the geometry of our experiments, the jet-like fluid movement is transformed into a steady fluid streaming which forms vortexes in the vicinity of the vibrating body even at low frequencies (Costalonga et al., 2015). The vortexes can facilitate fluid mixing close to the vibrating body, which is the inner surface of the vibrating membrane in our experiments. Thus, in the fluorescent dye experiments, this mixing should change the boundary condition at the closed pipe end and lead to additional spread of the diffusion front without changing its dispersion (Figure 5D). The diffusion front dispersion over the same time is larger for substances with larger diffusion coefficients. Therefore, we can predict that the effective range of increased concentration should be larger for salicylate used in our experiments (salicylate diffusion coefficient is 9.59×10^−4^ mm^2^/s (Lide (2002)) and for dexamethasone, the most frequently used drug for intratympanic treatment of hearing disorders (dexamethasone diffusion coefficient calculated from Stokes-Einstein equation which, however, underestimates experimental values is 6.82×10^−4^ mm^2^/s), than we observed for Lucifer yellow (diffusion coefficient is 3.1×10^−4^ mm^2^/s (Brink & Ramanan, 1985)).

The second major difference between our *in vivo* and fluorescent dye experiments is in the amount of material available for diffusion. The amount of dye was limited by its initial injection. A relatively large volume of 5 μl of salicylate solution was placed on the outer surface of the RW *in vivo*. Hence, an additional amount of salicylate could enter the ST down its concentration gradient when the salicylate concentration in the immediate vicinity of the inner surface of the RW dropped due to enhanced mixing because of the vortex formation described above and because the RW permeability increased during its mechanical stimulation (e.g. Park & Moon, 2014; Liao et al., 2020). This facilitated additional influx of salicylate could increase its concentration at the cochlear base, which is indicated by higher basal CAP threshold elevations observed in our experiments (Figure 4). The increase in salicylate concentration changes the diffusion boundary condition and promotes diffusion of salicylate to the cochlear apex. It should be noted that salicylate was utilized in this study due to its well documented physiological effects. However, it is a difficult drug to distribute along the cochlea because it is cleared rapidly from the ST (Sadreev et al., 2019). It is anticipated that drugs, which are better retained in the ST, will be redistributed along the cochlea even more quickly and efficiently (Salt and Ma, 2001; Sadreev et al., 2019).

This work is a proof of concept study and it remains to be demonstrated that the RW micro vibrations can promote distribution of substances for cochleae of the human cochlea’s size and for stimulation parameters that are safe for human cochlear function. If this drug delivery method is effective in human patients, it could be used to deliver and distribute drugs along the cochlea when cochlear pumping (Lukashkin et al., 2020) cannot be applied. For example, when the ossicular functionality is absent or impeded, e.g. after injection of high concentrations of hydrogel formulations into the middle ear (e.g. Piu et al., 2011; Schilder et al., 2019). Cochlear drug delivery utilizing micro vibrations of the RW could be particularly useful in patients with round window vibroplasty (e.g. Beltrame et al., 2014) if a part of the RW is left available for drug diffusion from the middle ear. In this case a vibrator is already present at the RW and any additional interventions required are minimal.

## Data availability

Data is available on request through the University of Brighton Research Data Repository at https://researchdata.brighton.ac.uk/

## References

Aris R. (1960). On the dispersion of a solute in pulsating flow through a tube. Proc R Soc A 259:370–376.

Beltrame AM, Todt I, Sprinzl G, Profant M, Schwab B. (2014). Consensus statement on round window vibroplasty. Ann Otol Rhinol Laryngol 123:734–740.

Borkholder DA, Zhu X, Frisina RD. (2014). Round window membrane intracochlear drug delivery enhanced by induced advection. J Control Release 174:171–176. doi: 10.1016/j.jconrel.2013.11.021

Brink PR, Ramanan SV. (1985). A model for the diffusion of fluorescent probes in the septate giant axon of earthworm. Axoplasmic diffusion and junctional membrane permeability. Biophys J 48:299–309.

Burwood GWS, Russell IJ, Lukashkin AN. (2017). Rippling pattern of distortion product otoacoustic emissions evoked by high-frequency primaries in guinea pigs. J Acoust Soc Am 142:855–862. doi:10.1121/1.4998584

Cheatham MA, Naik K, Dallos P. (2011). Using the cochlear microphonic as a tool to evaluate cochlear function in mouse models of hearing. J Assoc Res Otolaryngol 12:113–125.

Costalonga M, Brunet P, Peerhossaini H. (2015). Low frequency vibration induced streaming in a Hele-Shaw cell. Phys Fluids 27:013101. doi:10.1063/1.4905031

Creber NJ, Eastwood HT, Hampson AJ, Tan J, O’Leary SJ. (2018). A comparison of cochlear distribution and glucocorticoid receptor activation in local and systemic dexamethasone drug delivery regimes. Hear Res 368:75–85. doi: 10.1016/j.heares.2018.03.018

Dasgupta S. (2015). An Experimental Study of Dispersion in Oscillating Flows in Cylindrical Tubes. Thesis. Rochester Institute of Technology. Accessed from https://scholarworks.rit.edu/theses/8762

El Kechai N, Agnely F, Mamelle E, Nguyen Y, Ferrary E, Bochot A. (2015). Recent advances in local drug delivery to the inner ear. Int J Pharm 494:83–101. doi: 10.1016/j.ijpharm.2015.08.015

Ghiz AF, Salt AN, DeMott JE, Henson MM, Henson Jr OW, Gewalt SL. (2001). Quantitative anatomy of the round window and cochlear aqueduct in guinea pigs. Hear Res 162:105–112. doi: 10.1016/S0378-5955(01)00375-6

Greenwood DD. (1990). A cochlear frequency-position function for several species‐-29 years later. J Acoust Soc Am 87:2592–2605. doi: 10.1121/1.399052

Haghpanahi M, Gladstone MB, Zhu X, Frisina RD, Borkholder DA. (2013). Noninvasive technique for monitoring drug transport through the murine cochlea using micro-computed tomography. Ann Biomed Eng 41:2130–2142. doi: 10.1007/s10439-013-0816-4

Hallworth R. (1997). Modulation of outer hair cell compliance and force by agents that affect hearing. Hear Res 114:204–212.

Hao J, Li SK. (2019). Inner ear drug delivery: Recent advances, challenges, and perspective. Eur J Pharm Sci 126:82–92. doi: 10.1016/j.ejps.2018.05.020

Liao AH, Wang CH, Weng PY, Lin YC, Wang H, Chen HK, Liu HL, Chuang HC, Shih CP. (2020). Ultrasound-induced microbubble cavitation via a transcanal or transcranial approach facilitates inner ear drug delivery. JCI insight 5:e132880.

Lide DR. (2002). CRC Handbook of Chemistry and Physics, 83rd Edn. Boca Raton, FL: CRC Press.

Lukashkin AN, Sadreev II, Zakharova N, Russell IJ, Yarin YM. (2020). Local Drug Delivery to the Entire Cochlea without Breaching Its Boundaries. iScience 23:100945.

Marquardt T, Hensel J, Mrowinski D, Scholz G. (2007). Low-frequency characteristics of human and guinea pig cochleae. J Acoust Soc Am 121:3628–3638. https://doi.org/10.1121/1.2722506

Mynatt R, Hale SA, Gill RM, Plontke SK, Salt AN. (2006). Demonstration of a longitudinal concentration gradient along scala tympani by sequential sampling of perilymph from the cochlear apex. J Assoc Res Otolaryngol 7:182–193. doi: 10.1007/s10162-006-0034-y

Nyberg S, Abbott NJ, Shi X, Steyger PS, Dabdoub A. (2019). Delivery of therapeutics to the inner ear: The challenge of the blood-labyrinth barrier. Sci Transl Med 11:eaao0935.

Oliver D, He DZ, Klocker N, Ludwig J, Schulte U, Waldegger S, Ruppersberg JP, Dallos P, Fakler B. (2001). Intracellular anions as the voltage sensor of prestin, the outer hair cell motor protein. Science 292:2340–2343.

Park SH, Moon IS. (2014). Round window membrane vibration may increase the effect of intratympanic dexamethasone injection. Laryngoscope 124:1444–1451.

Patuzzi RB, Yates GK, Johnstone BM. (1989). Outer hair cell receptor current and sensorineural hearing loss. Hear Res 42:47–72.

Piu F, Wang X, Fernandez R, Dellamary L, Harrop A, Ye Q, Sweet J, Tapp R, Dolan DF, Altschuler RA, Lichter J. (2011). OTO-104: a sustained-release dexamethasone hydrogel for the treatment of otic disorders. Otol Neurotol 32:171–179.

Plontke SK, Biegner T, Kammerer B, Delabar U, Salt AN. (2008). Dexamethasone concentration gradients along scala tympani after application to the round window membrane. Otol Neurotol 29:401–406. doi: 10.1097/MAO.0b013e318161aaae

Plontke SK, Mynatt R, Gill RM, Borgmann S, Salt AN. (2007). Concentration gradient along the scala tympani after local application of gentamicin to the round window membrane. Laryngoscope 117:1191–1198. doi: 10.1097/MLG.0b013e318058a06b

Rivera T, Sanz L, Camarero G, Varela-Nieto I. (2012). Drug delivery to the inner ear: strategies and their therapeutic implications for sensorineural hearing loss. Curr Drug Deliv 9:231–242. doi: 10.2174/156720112800389098

Russell IJ, Schauz C. (1995). Salicylate ototoxicity: Effects on stiffness and electromotility of outer hair cells isolated from the guinea pig cochlea. Auditory Neurosci 1:309–319.

Sadreev II, Burwood GWS, Flaherty SM, Kim J, Russell IJ, Abdullin TI, Lukashkin AN. (2019). Drug diffusion along an intact mammalian cochlea. Front Cell Neurosci 13:161.

Salt AN, Lichtenhan JT, Gill RM, Hartsock JJ. (2013). Large endolymphatic potentials from low-frequency and infrasonic tones in the guinea pig. J Acoust Soc Am 133:1561–1571. doi: 10.1121/1.4789005

Salt AN, Ma Y. (2001). Quantification of solute entry into cochlear perilymph through the round window membrane. Hear Res 154:88–97. doi: 10.1016/S0378-5955(01)00223-4

Salt AN, Plontke SK. (2009). Principles of local drug delivery to the inner ear. Audiol Neurotol 14:350–360. doi: 10.1159/000241892

Schilder AGM, Su MP, Blackshaw LL, Staecker H, Lenarz T, Safieddine S, Gomes-Santos CS, Holme R, Warnecke A. (2019). Hearing protection, restoration, and regeneration: an overview of emerging therapeutics for inner ear and central hearing disorders Otol Neurotol 40:559–570

Shokrian M, Knox C, Kelley DH, Nam JH. (2020). Mechanically facilitated micro-fluid mixing in the organ of Corti. Sci Rep 10:14847. doi: 10.1038/s41598-020-71380-5

Taylor GI. (1953). Dispersion of soluble matter in solvent flowing slowly through a tube. Proc R Soc A 219:186–203.

Thorne M, Salt AN, DeMott JE, Henson MM, Henson Jr OW and Gewalt SL. (1999). Cochlear fluid space dimensions for six species derived from reconstructions of three-dimensional magnetic resonance images. Laryngoscope 109:1661–1668.

Watson EJ. (1983). Diffusion in oscillatory pipe flow. J Fluid Mech 133:233–244.

Weddell TD, Yarin YM, Drexl M, Russell IJ, Elliott SJ, Lukashkin AN. (2014). A novel mechanism of cochlear excitation during simultaneous stimulation and pressure relief through the round window. J R Soc Interface 11:20131120. doi: 10.1098/rsif.2013.1120

Wysocki J, Sharifi M. (2005). Measurements of selected parameters of the guinea pig temporal bone. Folia Morphol 64:145–150.

